# Graph Convolutional Learning of Multimodal Brain Connectome Data for Schizophrenia Classification

**DOI:** 10.1101/2023.01.05.522960

**Authors:** Sanjay Ghosh, Eshan Bhargava, Chieh-Te Lin, Srikantan S. Nagarajan

## Abstract

The long term goal of this work is to develop powerful tools for brain network analysis in order to study structural and functional connectivity abnormalities in psychiatric disorders like schizophrenia. Graph convolutional neural networks (GCNN) are quite effective for learning complex discriminate features in graph-structured data. Here, we explore the GCNN to learn the discriminating features in multimodal human brain connectomes for the purpose of schizophrenia disorder classification. In particular, we train and validate a network using both structural connectivity graphs obtained from diffusion tensor imaging data and functional connectivity from functional magnetic resonance imaging data.We compare the GCNN method with a support vector machine based classifier and other popular classification benchmarks. We demonstrate that the proposed graph convolution method has the best performance compared to existing benchmarks with F1 scores of 0.75 for schizophrenia classification. This demonstrates the potential of this approach for multimodal diagnosis and prognosis in mental health disorders.

## 1. INTRODUCTION

Schizophrenia is a severe psychiatric disorder which causes impairments in memory, attention, and other high-order cognitive dysfunctions. Recent advances in magnetic resonance imaging (MRI) have made it possible to examine the changes in white and grey matter in the brain of patients suffering from schizophrenia. Various functional neuroimaging techniques are also being used to understand the abnormalities in neural activities of schizophrenia patients. A recent promising research direction in context of neuroimaging analysis is the conceptualization of a particular disorder as a dysconnectivity syndrome [1]. Experimental studies have shown the evidence of alterations in both structural and functional brain connectivity of schizophrenia patients [2, 3]. In fact, a thorough characterization of the structural and functional connectome is of importance to the development of novel biomarkers both for prediction and treatment of schizophrenia [4]. Having such biomarkers in schizophrenia could lead to clinically useful tools for establishing both diagnosis and prognosis of the disease. From a methodology perspective, the choice of biomarkers can be addressed as a feature selection problem. The aim is to capture relevant features enabling us to differentiate patients from control subjects accurately.

Deep learning methods have been recently used to analyze the whole brain connectome or neuroimaging data for various brain disorders analysis [5, 6, 7, 8]. Ktena et al [5] proposed a graph neural network to analyze brain functional connectivity networks for autism classification. A multi-scale graph convolutional network (MMTGCN) was introduced for brain connectivity based analysis. In particular, it was employed for classification of multiple disorders attention-deficit/hyperactivity disorder (ADHD), mild cognitive impairment (MCI) and cerebral small vessel disease (cSVD). Zhang et al. [9] proposed a graph neural network using multiple modalities of brain images in relationship prediction which is useful for distinguishing Parkinson’s disease (PD) cases from controls. A multiclass classifier for classification of subjects on the Alzheimer’s disease (AD) spectrum using structural connectivity graphs based on diffusion tensor imaging (DTI) in [1]. We note that not much effort has been made in machine learning based classification of schizophrenia disorder. Authors in [10] proposed a support vector classifier (SVC) based classifier that aims to classify brain connectomes by maximizing the margin between classes of healthy subjects and schizophrenia patients. They also investigated the performance of the classifier for connectomes with different resolutions.

In this paper, we propose a graph convolutional neural network which learns to distinguish between brain networks of schizophrenia patients and healthy controls. From a conceptual view of graph network, we use a structural connectome as the underlying graph structure and the concerned input graph signal is derived from the functional connectomes. In particular, the graph signal at each node is the vector containing functional similarity to rest of the nodes in brain.

This paper is organized as follows. In Section 2, we introduce the basics of the GCNN, it’s implementation, and the steps of dataset preparation. In Section 3, we present the results of schizophrenia classification performance along with the details of quality metrics used for evaluation. Finally, we conclude our findings in Section 4.

## 2. METHOD

### 2.1. Graph Convolution on Brain Network

Human brain networks are known to depict activity patterns within the brain, and these connectivity patterns have a unique graph architectures at the whole brain scale. Nodes in a graph can represent region-of-interests (ROIs) accompanied by a set of features. To learn the topological attributes of both brain structural and functional connectivity networks, we explore the unique nature of graph convolutional neural networks. In particular, the idea is to learn the functional characteristics of the brain via convolutional deep network defined on the structural graph. Visual examples of cohort level connectomes are shown in Figs. 1 and 2. Notice the subtle differences between each pairs. All four connectomes are from a public dataset [11]. Each connectome has 83 ROIs out of which 68 ROIs are in cortical and rest 15 ROIs are in sub-cortical regions of the brain.

**Fig. 1.**
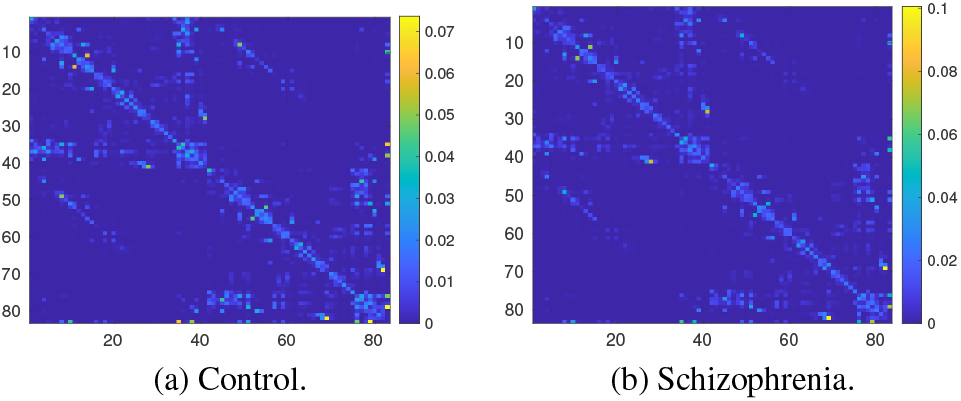
Cohort-level structural (DTI) connectomes.

**Fig. 2.**
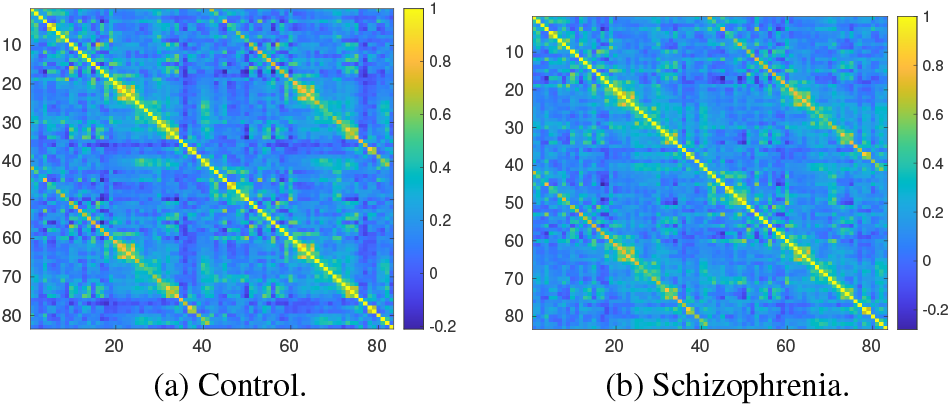
Cohort-level functional (fMRI) connectomes.

We construct the graph 𝒢 from structural connectome ***S***; where the number of nodes *N* is the number of ROIs in the connectome. Each edge weight in 𝒢 is given by the connection strength between a pair of ROIs. The normalized Laplacian is defined as:

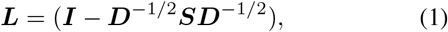

where the degree matrix *D* = diag_*i*_ [Σ_*i*_ ***S***_*ij*_] is obtained from the structural connectome. Graph signal at each node is derived from the functional connectome ***F*** = [*f*_1_, *f*_2_, …, *f*_*N*_]^*T*^. In other words, the input graph signal at node *i* is given by *x*_*i*_ = *f*_*i*_, ∀*i* = {1, 2, …, *N*}. The spectral convolution at *k*^*th*^ layer is defined on graph 𝒢 as follows:

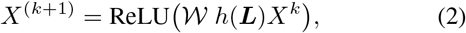

where 𝒲 is a learnable weight matrix of the layer and *h* is arbitrary function. Suppose *h*(***L***) = *Uh*(Λ)*U*^*T*^, where *U* is the matrix whose columns are eigenvectors and Λ is a diagonal matrix with eigenvalues of ***L***. For the sake of completeness, the input graph signal at the 0^*th*^ layer is given by *X*^0^ = [*f*_1_, *f*_2_, …, *f*_*N*_]^*T*^. To overcome the computational bottleneck of computing the eigenvectors of ***L***, authors in [12], proposed Chebyshev polynomial approximation of the spectral filtering:

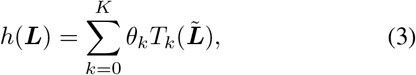

where 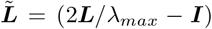. Furthermore, the Chebyshev polynomial can be computed recursively. Therefore, there is no need of computing the eigen decomposition. Finally, the trainable variables are weight matrix 𝒲 and Chebyshev coefficients vector *θ* of length (*K* + 1).

### 2.2. Proposed Deep Architecture

Our proposed deep network is shown in Fig. 3. As the theme of research suggests, the core component of the proposed architecture is a graph convolutional neural network is in (2). The network contained five layers including three graph convolutional layers (denoted GCN) and two fully connected layers, each followed by a rectified linear unit (denoted ReLU). Finally we have a softmax output layer computing class-membership probabilities. We use cross-entropy loss to guide the learning of the proposed classification network. Note that our deep network is relatively low-weighted. In fact, we refer [13] which reported a study the effect of numbers of graph convolutional layers in autism prediction.

**Fig. 3.**
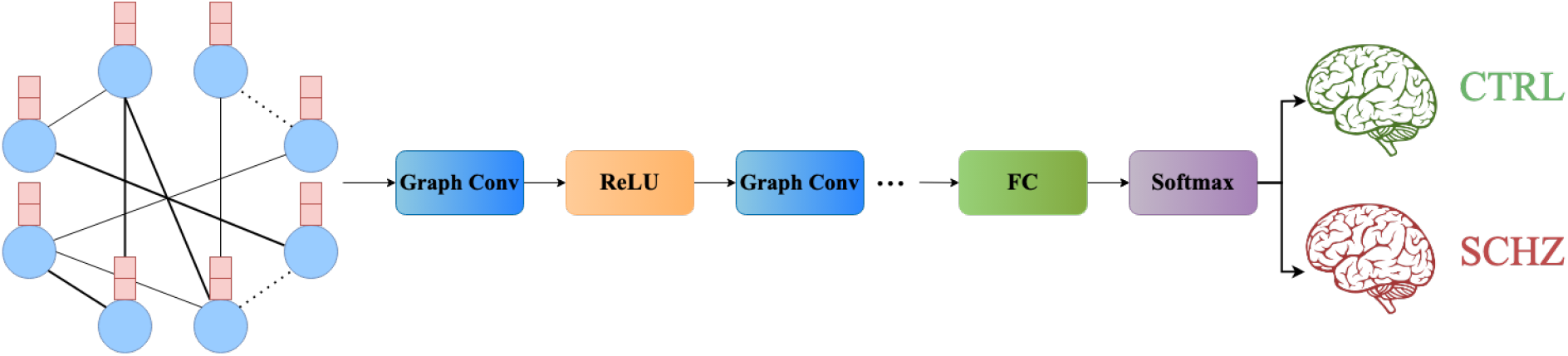
Architecture of our deep learning model to classify schizophrenia (SCHZ) patients and healthy control (CTRL) subjects. The step of multimodal brain connectome integration is also demonstrated using color-coding. Notice that the graph signal (light brown) in the input comes from functional brain connectome. On the other hand, the graph structure (light blue) i.e. edge weights are constructed from structural connectome data.

### 2.3. Data Augmentation

Availability of sufficient amount of data to adequately train a deep network is often a real concern in brain image analysis. Data augmentation is a process to create new artificial data by inducing artificial perturbation to the available data set. It is known in the literature to be effective for increasing robustness and even performance boosting. Driven by the symmetric nature of brain connectomes, we propose to add random Gaussian noise (symmetric) as follows:

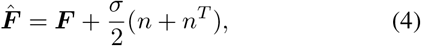

where *n* is a *N × N* matrix whose entries are drawn from 𝒩(0, 1). We basically perform augmentation only on the functional data. However, no perturbation is performed on the graph structure.

### 2.4. Dataset

The dataset used in our experiments are downloaded from [11]. The cohort consists of a schizophrenic group of 27 subjects and a control group of 27 healthy subjects. All of the data was collected at the Service of General Psychiatry at the Lausanne University Hospital. The schizophrenic patients were diagnosed with schizophrenic and schizo-affective disorders after meeting the DSM-IV criteria. Control subjects had no history of neurological disease. All 54 subjects had given written consent following the institutional guidelines approved by the Ethics Committee of Clinical Research of the Faculty of Biology and Medicine, University of Lausanne, Switzerland.

### 2.5. Data Preparation

A 3 Tesla Siemens Trio scanner with a 32-channel head coil was used for scanning all subjects. The data collection protocol consisted of three imaging modalities: (A) a magnetization-prepared rapid acquisition gradient echo (MPRAGE) sequence sensitive to white/gray matter contrast (1-mm in-plane resolution, 1.2-mm slice thickness), (B) a diffusion spectrum imaging (DSI) sequence (128 diffusion-weighted volumes and a single b0 volume, maximum b-value 8,000 s/mm2, 2.2×2.2×3.0 mm voxel size), and (B) a gradient echo planar imaging (EPI) sequence sensitive to BOLD contrast (3.3-mm in-plane resolution and slice thickness with a 0.3-mm gap, TE 30 ms, TR 1,920 ms, resulting in 280 images per participant).

Structural connectivity matrices were computed using deterministic streamline tractography on reconstructed DSI data. The pipeline was initiated with 32 streamline propagations per diffusion direction, per white matter voxel. Structural connectivity between pairs of regions was measured in terms of fiber density between each pair of ROIs. We note that fiber density is defined as the number of streamlines between the two regions, normalized by the average length of the streamlines and average surface area of the two regions. Functional connectomes were estimated from fMRI BOLD time-series. In particular, the absolute value of the Pearson correlation was computed between individual brain regions’ time-courses.

## 3. RESULT

### 3.1. Evaluation Metric and Benchmarks

We examine the effectiveness of our multimodal classification method on a public dataset of schizophrenia [11]. We perform 3-fold cross validation in the training process. Specifically, we randomly select 1/3 of the sample size from each class as the testing dataset, while the remaining 2/3 of the subjects are treated as the training dataset. This way we successfully utilize the existing available subjects to produce an unbiased performance. To evaluate the classification performance, We use six different metrics as follows: accuracy (ACC), sensitivity (SENS), specificity (SPEC), positive predictive values (PPV), negative predictive values (NPV) and *F*_1_ score.

We compare the classification performance of our method with three conventional learning-based methods: K-nearest neighbour (KNN), logistic regression, and K-means clustering. Finally, we compare with the state-of-the-art method [10] for schizophrenia in the literature.

### 3.2. Classification Performance

Our deep learning model was trained and tested using the TensorFlow python platform. During training, we use crossentropy loss function which was minimized using the stochastic gradient descent (SGD) with momentum algorithm. The algorithm parameters are as follows: learning rate 0.001, batch size 18, decay rate 0.998, momentum 0.95 and 800 training epochs.

We start with baseline and competing approaches to evaluate the classification performance. For all six quality metrics, a higher value indicates better classification performance. We present the summary of our experimental results in Table 1.

**Table 1.**
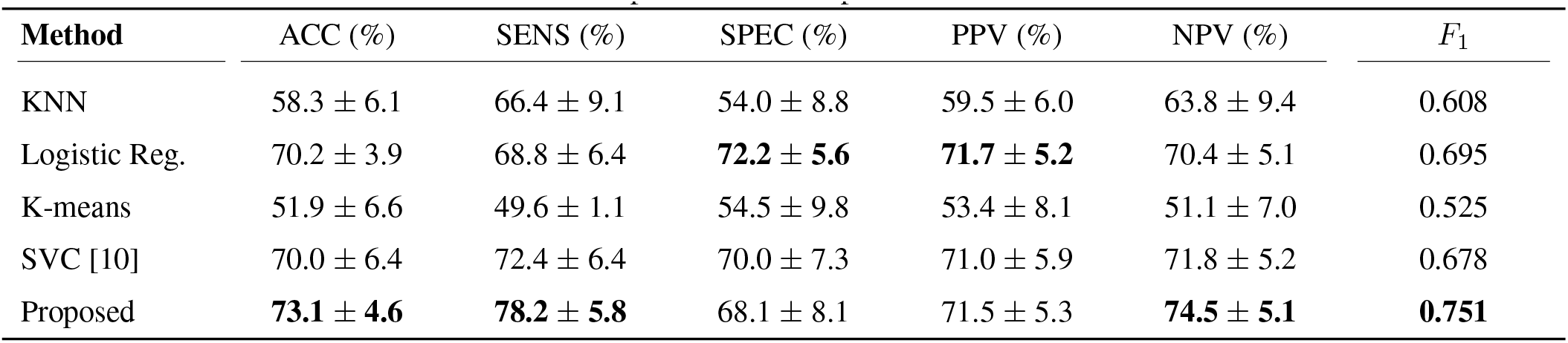
Comparison of schizophrenia classification.

| Method | ACC (%) | SENS (%) | SPEC (%) | PPV (%) | NPV (%) | $F_1$ |
| --- | --- | --- | --- | --- | --- | --- |
| KNN | 58.3 ± 6.1 | 66.4 ± 9.1 | 54.0 ± 8.8 | 59.5 ± 6.0 | 63.8 ± 9.4 | 0.608 |
| Logistic Reg. | 70.2 ± 3.9 | 68.8 ± 6.4 | **72.2 ± 5.6** | **71.7 ± 5.2** | 70.4 ± 5.1 | 0.695 |
| K-means | 51.9 ± 6.6 | 49.6 ± 1.1 | 54.5 ± 9.8 | 53.4 ± 8.1 | 51.1 ± 7.0 | 0.525 |
| SVC [10] | 70.0 ± 6.4 | 72.4 ± 6.4 | 70.0 ± 7.3 | 71.0 ± 5.9 | 71.8 ± 5.2 | 0.678 |
| Proposed | **73.1 ± 4.6** | **78.2 ± 5.8** | 68.1 ± 8.1 | 71.5 ± 5.3 | **74.5 ± 5.1** | **0.751** |

We report the mean and standard deviation of each entry after performing 30 realizations. It is evident that our proposed method offers the highest classification accuracy when compared with the other four methods. Notice that the *F*_1_ score obtained by our method is significantly better. We highlight the improvement over the SVC based method [10] designed for schizophrenia analysis.

## 4. CONCLUSION

In this paper, we proposed a graph neural network method from multimodal human connectome to develop a robust biomarker for the identification of subjects with schizophrenia. As inherent in deep learning, we perform an automatic feature selection from the input aiming to retrieve more meaningful biomarkers performing accurately on the identification of schizophrenia versus healthy controls. By efficiently integrating both structural and functional connectivity matrices as a multi-modal representation of connectomes, our method was able to achieve more accurate schizophrenia classification. In future research, we plan to further improve our method by combining multiscaled higher resolution connectomes within our deep model. In a different direction, we plan to explore deeper understanding of clinical relevance. This would also include addressing the question whether the alteration of structural or functional connectomes play a more important role for the task of schizophrenia classification.

## Acknowledgment

The authors would like to thank Patric Hagmann for publicly sharing the dataset at Connectomics Lab webpage.

## REFERENCES

[1] T.-A. Song, S. R. Chowdhury, F. Yang, H. Jacobs, G. El Fakhri, Q. Li, K. Johnson, and J. Dutta, “Graph convolutional neural networks for alzheimer’s disease classification,” IEEE 16th International Symposium on Biomedical Imaging (ISBI 2019), pp. 414–417, 2019.

[2] M.-E. Lynall, D. S. Bassett, R. Kerwin, P. J. McKenna, M. Kitzbichler, U. Muller, and E. Bullmore, “Functional connectivity and brain networks in schizophrenia,” Journal of Neuroscience, vol. 30, no. 28, pp. 9477–9487, 2010.

[3] A. Fornito, A. Zalesky, C. Pantelis, and E. T. Bullmore, “Schizophrenia, neuroimaging and connectomics,” Neuroimage, vol. 62, no. 4, pp. 2296–2314, 2012.

[4] D. I. Kim, J. Sui, S. Rachakonda, T. White, D. S. Manoach, V. P. Clark, B.-C. Ho, S. C. Schulz, and V. D. Calhoun, “Identification of imaging biomarkers in schizophrenia: a coefficient-constrained independent component analysis of the mind multi-site schizophrenia study,” Neuroinformatics, vol. 8, no. 4, pp. 213–229, 2010.

[5] S. I. Ktena, S. Parisot, E. Ferrante, M. Rajchl, M. Lee, B. Glocker, and D. Rueckert, “Metric learning with spectral graph convolutions on brain connectivity networks,” NeuroImage, vol. 169, pp. 431–442, 2018.

[6] G. Ma, N. K. Ahmed, T. L. Willke, D. Sengupta, M. W. Cole, N. B. Turk-Browne, and P. S. Yu, “Deep graph similarity learning for brain data analysis,” in Proceedings of the 28th ACM International Conference on Information and Knowledge Management, 2019, pp. 2743–2751.

[7] C. J. Brown, J. Kawahara, and G. Hamarneh, “Connectome priors in deep neural networks to predict autism,” in 2018 IEEE 15th international symposium on biomedical imaging (ISBI 2018). IEEE, 2018, pp. 110–113.

[8] C.-T. Lin, S. Ghosh, L. B. Hinkley, C. L. Dale, A. C. Souza, J. H. Sabes, C. P. Hess, M. E. Adams, S. W. Cheung, and S. S. Nagarajan, “Multi-tasking deep network for tinnitus classification and severity prediction from multimodal structural mr images,” Journal of Neural Engineering, 2022.

[9] X. Zhang, L. He, K. Chen, Y. Luo, J. Zhou, and F. Wang, “Multi-view graph convolutional network and its applications on neuroimage analysis for parkinson’s disease,” in AMIA Annual Symposium Proceedings, vol. 2018. American Medical Informatics Association, 2018, p. 1147.

[10] L. Gutiérrez-Gómez, J. Vohryzek, B. ChiAm, P. S. Baumann, P. Conus, K. Do Cuenod, P. Hagmann, and J.-C. Delvenne, “Stable biomarker identification for predicting schizophrenia in the human connectome,” NeuroImage: Clinical, vol. 27, p. 102316, 2020.

[11] J. Vohryzek, Y. Aleman-Gomez, A. Griffa, J. Raoul, M. Cleusix, P. S. Baumann, P. Conus, K. D. Cuenod, and P. Hagmann, “Structural and functional connectomes from 27 schizophrenic patients and 27 matched healthy adults [data set],” Zenodo, 2020.

[12] M. Defferrard, X. Bresson, and P. Vandergheynst, “Convolutional neural networks on graphs with fast localized spectral filtering,” Advances in neural information processing systems, vol. 29, 2016.

[13] Y. Ma, D. Yan, C. Long, D. Rangaprakash, and G. Deshpande, “Predicting autism spectrum disorder from brain imaging data by graph convolutional network,” International Joint Conference on Neural Networks (IJCNN), pp. 1–8, 2021.

